# Sequential Multiple Assignment Randomized Trials: An Opportunity for Improved Design of Stroke Reperfusion Trials

**DOI:** 10.1101/035683

**Authors:** William Meurer, Nicholas J. Seewald, Kelley Kidwell

## Abstract

Background: Modern clinical trials in stroke reperfusion fall into two categories: alternative systemic pharmacological regimens to alteplase and "rescue" endovascular approaches using targeted thrombectomy devices and/or medications delivered directly for persistently vessel occlusions. Clinical trials in stroke have not evaluated how initial pharmacological thrombolytic management might influence subsequent rescue strategy. A sequential multiple assignment randomized trial (SMART) is a novel trial design that can test these dynamic treatment regimens and lead to treatment guidelines which more closely mimic practice.

Aim: To characterize a SMART design in comparison to traditional approaches for stroke reperfusion trials.

Methods: We conducted a numerical simulation study that evaluated the performance of contrasting acute stroke clinical trial designs of both initial reperfusion and rescue therapy. We compare a SMART design where the same patients are followed through initial reperfusion and rescue therapy within one trial to a standard phase III design comparing two reperfusion treatments and a separate phase II futility design of rescue therapy in terms of sample size, power, and ability to address particular research questions.

Results: Traditional trial designs can be well powered and have optimal design characteristics for independent treatment effects. When treatments, such as the reperfusion and rescue therapies, may interact, commonly used designs fail to detect this. A SMART design, with similar sample size to standard designs, can detect treatment interactions.

Conclusions: The use of SMART designs to investigate effective and realistic dynamic treatment regimens is a promising way to accelerate the discovery of new, effective treatments for stroke.

## Main Manuscript

### Introduction

Since 1995, intravenous alteplase is the only pharmacological treatment approved for use to reduce disability following acute ischemic stroke.(1) Despite its lone position in the stroke pharmacological arsenal, a large proportion of treated patients do not improve. Two primary reasons likely drive this, namely failure to re-open the affected blood vessel or restoration of blood flow that occurs after permanent injury to brain tissue. In carefully selected patients, endovascular treatments after intravenous thrombolysis can remarkably improve patient outcomes.

Modern clinical trials in stroke reperfusion have been broadly grouped into two categories: alternative systemic pharmacological regimens to alteplase and "rescue" endovascular approaches using targeted thrombectomy devices and/or medications delivered directly to the persistently occluded blood vessel. Several recent trials that predominately included patients treated with alteplase using a variety of selection criteria demonstrated that mechanical thrombectomy improved clinical outcomes.(2-4) Clinical trial design in this field has not frequently evaluated the interplay between initial pharmacological thrombolytic management and subsequent rescue strategy. In addition, endovascular treatments are available in a small portion of hospitals worldwide. Ischemic stroke is a time-sensitive disease, and patients need rapid treatment. The current clinical trial paradigm cannot adequately answer important treatment questions. For example, if a new pharmacological regimen is superior to alteplase, will endovascular approaches have the same, better, or worse efficacy and safety? In addition, might faster pharmacological alternatives or adjuncts to endovascular approaches such as rescue doses of alternative thrombolytics or other adjunctive treatments also be safe and effective and expand the number of hospitals that can rapidly address initial alteplase (or new regimen) failures in patients with large vessel occlusions.

Newer methods in clinical trials likely can improve discovery in this arena. While clinicians use sequentially accruing information to make decisions at the bedside, this information is infrequently incorporated into clinical trials. A dynamic treatment regimen (DTR) is a guideline specifying a sequence of treatments based on individual characteristics, behaviors, or response.(5) A DTR includes an initial treatment, an intermediate outcome, and subsequent treatment based on each intermediate outcome option. For example, one DTR for the treatment of stroke is to first treat the patient with alteplase. If the patient has an NIHSS score ≤ 7 after two hours from alteplase administration, continue to monitor patient; if the patient has an NIHSS >7 after two hours from alteplase administration, take the patient to endovascular intervention. Therefore, a DTR is a tool for physicians for which to base treatment for patients at the onset of disease and to continue throughout time adjusting to the patient. DTRs formalize the way physicians practice medicine depending on patient outcomes throughout time, but appear to patients to be just the sequence of treatments that they receive. In order to construct and provide evidence for beneficial or optimal DTRs, we need to extend the standard (one-stage) randomized control trial design.

This extension to construct and compare DTRs is called a sequential multiple assignment randomized trial.(6, 7) This type of trial follows the same patients throughout two or more stages, re-randomizing patients to subsequent therapy based on intermediate outcomes. The trial is similar to a sequential factorial design where randomization at later stages depends on patient characteristics (e.g. response to treatment at previous stage). The goal of a SMART is to inform the development of DTRs and to potentially find an optimal DTR that will more closely mimic the treatment process.

## Aims and Objectives

Our objective is to demonstrate the utility of an alternate design strategy (SMART) in stroke reperfusion trials, an area where quick, sequential, individualized decisions are necessary and to compare this strategy to clinical trial designs traditionally used in stroke.

## Methods

### Overview

We conducted a numerical simulation study that evaluated the performance of contrasting acute stroke clinical trial designs of both initial reperfusion and rescue therapy. We used several scenarios for the "true" treatment effects of the various strategies. We evaluated two initial reperfusion strategies, which represent alteplase versus a new pharmacologic regimen. In addition, we evaluated two rescue therapies, which represent the current best endovascular approach versus rescue pharmacotherapy.

The scenarios reflected a range of possible situations, including for example, rescue pharmacotherapy only being effective if linked with the alteplase initial re-perfusion treatment. We evaluated the approach of doing the initial reperfusion and rescue trials separately, versus the SMART approach. The main outcome for any simulated trial was the proportion of true positive and false positive trials given the scenario in question over a range of plausible sample sizes.

### Design of Trial

An example of a SMART in the treatment of stroke is shown in Figure 1. Here there are two stages, one for the initial randomization to receive two distinct thrombolytic regimens, lytic A versus lytic B (i.e. A is alteplase and B is tenecteplase; or A is alteplase and B is the combination of alteplase and an additional agent such as a glycoprotein IIb/IIIa inhibitor), and then another stage based on response to NIHSS after two hours from lytic administration. The example is not specific to comparisons of single agents (e.g. first-stage treatments could include a single agent, A is alteplase, versus a combination, B is alteplase combined with eptifibitide). At 2 hours, responders are defined as those with NIHSS≤ 7 and these patients are monitored to make sure their response is stable (it is possible to have a SMART design where these patients are also re-randomized to a set of treatment options). Non-responders are those with an NIHSS>7 and are re-randomized to undergo mechanical thrombectomy versus a new adjunctive medication. The outcome of the trial is a categorization of overall response at 90 days based on the modified Rankin scale (mRS; success is defined for those who initially respond to the first-stage treatment as an mRS at 90 days of 0 or 1 and success is defined for those who do not initially respond and received follow-up treatment as an mRS of 0, 1 or 2).

**Figure 1:** Overview of a clinical trial incorporating a dynamic treatment regime. Caption: Cath Lab denotes when patients undergo mechanical thrombectomy.

There are four embedded DTRs in the SMART from Figure 1. Two DTRs begin with lytic A ([1] First give patient lytic A. If NIHSS≤ 7 then continue to monitor patient (usual care). If NIHSS≤7 then proceed for the patient to undergo mechanical thrombectomy (usual care); [2] First give patient lytic A. If NIHSS≤ 7 then continue to monitor patient. If NIHSS>7 then give the new adjunctive treatment). Two DTRs begin with lytic B ([1] First give patient lytic B.I If NIHSS≤ 7 then continue to monitor patient (usual care). If NIHSS>7 then proceed for the patient to undergo mechanical thrombectomy (usual care); [2] First give patient lytic B. If NIHSS≤ 7 then continue to monitor patient. If NIHSS>7 then give the new adjunctive treatment). Note that in Figure 1, each branch of treatment is not its own DTR; rather, a DTR recommends treatments for both responders and non-responders, and so is composed of two branches of treatment. DTRs that begin with the same treatment overlap by including one of the same treatment branches (here monitoring responder patients). In addition, while an NIHSS of 7 was used for this example – an alternate, rapidly available clinical parameter (e.g. presence of a large vessel occlusion) could easily be substituted for this as the information that provides the branch point.

### Base case scenarios

In clinical trial simulation, design performance is evaluated under a number of simulated truths. The scenarios that the SMART and the conventional clinical trial were simulated under are summarized in Table 1. For example, in the null scenario patients who receive lytic A or lytic B and respond (NIHSS≤7) have 76% probability of a good outcome at 90 days; similarly patients who do not initially respond but received either treatment in the second stage (undergoing mechanical thrombectomy or current usual care versus new adjunctive medication) have 30% probability of a good outcome at 90 days. It is important to simulate a null scenario like this to determine how frequently the design leads to a false positive conclusion. Scenario 2 subjects the clinical trials to the situation that lytic A is superior to B (76% of early responders have a good 90-day outcome with A versus only 57% with B) for the first stage treatments, and there is a 10% absolute increase in the proportion of initially non-responding subjects with a good outcome at 90 days (30% undergoing mechanical thrombectomy or current standard of care and 40% with new adjunctive medication) which was not different whether or not lytic A or B was the initial treatment. In this scenario, fewer lytic A patients were initial non-responders. Other scenarios follow directly from the table.

**Table 1:**
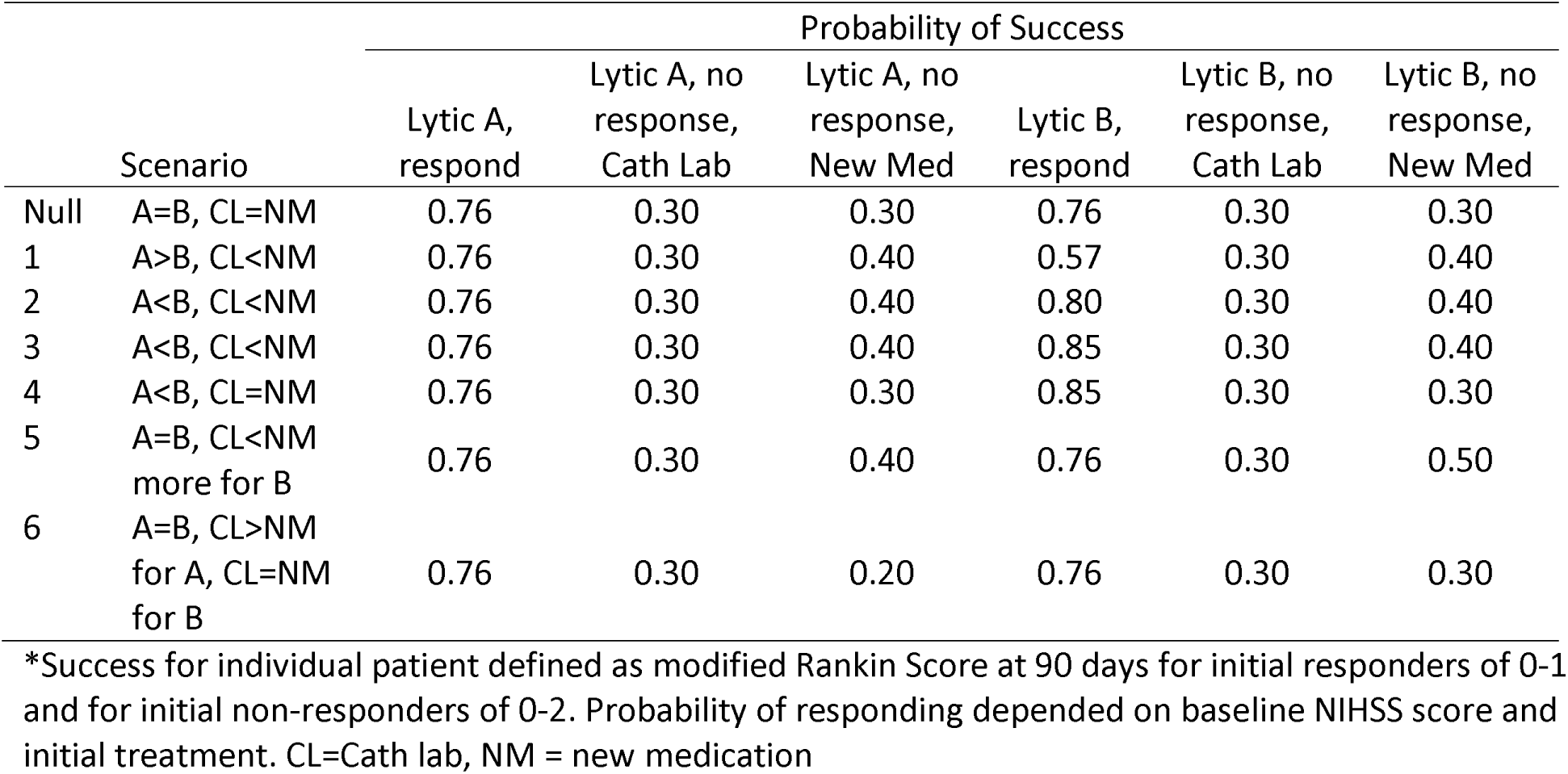
The overall probabilities of success for individuals following each treatment pathway.

|  |  | Probability of Success |  |  |  |  |  |
| --- | --- | --- | --- | --- | --- | --- | --- |
|  | Scenario | Lytic A, respond | Lytic A, no response, Cath Lab | Lytic A, no response, New Med | Lytic B, respond | Lytic B, no response, Cath Lab | Lytic B, no response, New Med |
| Null | A=B, CL=NM | 0.76 | 0.30 | 0.30 | 0.76 | 0.30 | 0.30 |
| 1 | A>B, CL<NM | 0.76 | 0.30 | 0.40 | 0.57 | 0.30 | 0.40 |
| 2 | A<B, CL<NM | 0.76 | 0.30 | 0.40 | 0.80 | 0.30 | 0.40 |
| 3 | A<B, CL<NM | 0.76 | 0.30 | 0.40 | 0.85 | 0.30 | 0.40 |
| 4 | A<B, CL=NM | 0.76 | 0.30 | 0.30 | 0.85 | 0.30 | 0.30 |
| 5 | A=B, CL<NM<br>more for B | 0.76 | 0.30 | 0.40 | 0.76 | 0.30 | 0.50 |
| 6 | A=B, CL>NM<br>for A, CL=NM<br>for B | 0.76 | 0.30 | 0.20 | 0.76 | 0.30 | 0.30 |

\*Success for individual patient defined as modified Rankin Score at 90 days for initial responders of 0-1 and for initial non-responders of 0-2. Probability of responding depended on baseline NIHSS score and initial treatment. CL=Cath lab, NM = new medication

### Simulation

We simulated the above-described SMART with sample sizes of 2000, 1500 and 700. In addition, we calculated the power for a traditional clinical trial comparing lytic A to lytic B under of the same sample sizes which are typically within the range of sample sizes used for phase III trials in acute stroke treatment (2000, 1500, 700). To mimic one of the favored designs for trial development in stroke, we assumed that a single arm futility trial (phase II) would first be used for the second stage treatment (new medicine versus undergoing mechanical thrombectomy in non-responders) and that this trial would draw from entirely different patients than the larger trial (phase III) comparing lytic A and B.(8) We assumed type I and type II errors of 0.1 for the futility trial and sample sizes of 200, 150 and 70. We also assumed that the historical rate of good outcome (mRS = 0 or 1) for this trial would be 30%. We simulated the trial to find the results of the futility trial (i.e. to go forward or declare futility) under three scenarios: the new medication having a 20% good outcome rate (harm), a 30% good outcome rate (null), or a 40% good outcome rate (benefit). Numerical simulations were performed with R version 3.1.3. We provide the code used the supplementary material.

## Results

### Performance of the traditional approach

The simulated phase III trial performed as expected – with a type I error of 0.05 (Table 2, power under the null case). High power was observed for scenario 1 (great superiority of A versus B). Scenarios 5 and 6 were similar to the null, as the differential effects in those scenarios were the different responses to the second-stage treatments (undergoing mechanical thrombectomy versus new adjunctive medication) which were not included in this trial. Scenario 2 had low power (relatively weak effect of B versus A), and scenarios 3 and 4 had reasonable power, even at a sample size of 700 (0.59).

**Table 2:**
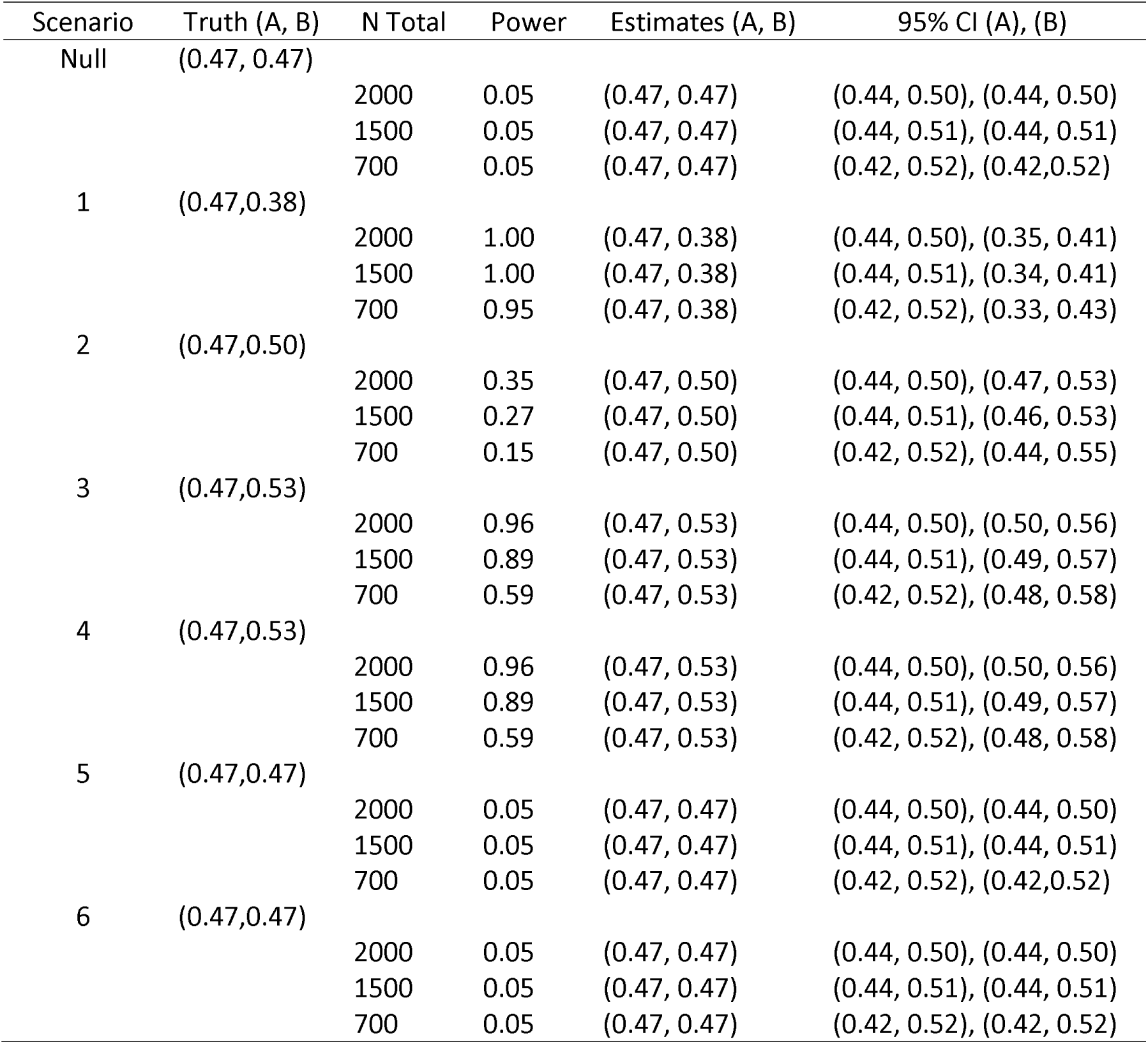
Sample Sizes, power, and estimated 95% confidence intervals (CIs) of treatment success for a standard phase III stroke trial comparing lytic A to lytic B

For the separate futility trial comparing the second stage treatments, we observed almost no chance of recommending taking the new treatment forward when it was harmful (Table 3), a relatively high likelihood (43.3%) of proceeding in the null case with low sample size (n=70), and a very high likelihood of proceeding with the new second stage treatment having a treatment effect size of 10% (about 95% for sample sizes of 70–200.)

**Table 3:** Results from a phase II single arm futility trial of new medicine following non-response to lytic Caption: The lower bound is the lower 90% confidence interval for 40% success (using the Wald method). The hypothesis for the futility trial is that there is a 0.1 increase in proportion of good outcomes from a baseline good outcome proportion of 0.3. For example, if the truth is that 30% of patients treated with the new medicine have a good outcome, and using a single arm futility trial of 70 patients to simulate this scenario, approximately 43% of trials will erroneously declare non-futility.

| Given<br>Probability of<br>Success for<br>2<sup>nd</sup>-stage | Sample Size | Lower<br>Bound to<br>continue | Proportion<br>successful<br>trials |
| --- | --- | --- | --- |
| 0.20 | 70 | 0.3037 | 0.018 |
|  | 150 | 0.3342 | 0 |
|  | 200 | 0.3430 | 0 |
| 0.30 | 70 | 0.3037 | 0.433 |
|  | 150 | 0.3342 | 0.164 |
|  | 200 | 0.3430 | 0.107 |
| 0.40 | 70 | 0.3037 | 0.947 |
|  | 150 | 0.3342 | 0.945 |
|  | 200 | 0.3430 | 0.953 |

### Performance of the SMART

In the null case for the trial of sample size 1500, each of the DTRs has estimated individual patient success probabilities of 0.471 or 0.472 (Table 4). This value represents the overall average good outcome rate within each of the groups, and takes into account responders (subjects who have a post first stage NIHSS ≤7) and non-responders who are eligible to receive a second stage treatment – new adjunctive medication versus usual care. Similar to the traditional case above, in scenario 1, comparing the DTRs that end with undergoing mechanical thrombectomy or usual care (lytic A or B followed with usual care) shows better outcomes with lytic A: 0.472 (95% CI 0.427-0.518) versus lytic B: 0.376 (95% 0.333-0.422). In addition, within Scenario 1, DTRs that end with the new medication are numerically better in terms of the probability of a good outcome than the DTRs that end with usual care in the second stage. Scenario 2 shows similar results for a small effect of DTRs that begin with new lytic B having better outcome than DTRs that begin with lytic A; again with DTRs that begin with new medicine in the second stage being better than DTRs that end with usual care. Scenario 3 is similar to scenario 2, but gives greater differences between DTRs that begin with lytic B vs. A and greater effect of DTRs that end with new medicine vs. usual care. Scenario 4 sets DTRs that begin with lytic A to have the same probability of successful outcome and DTRs that begin with lytic B to have a different from A, but equal to each other probability of a successful outcome such that the effect of second-stage treatment is equal across lytics. We simulated the DTR with a total sample size of 2000, and as expected the confidence intervals were slightly narrower (Supplemental Table 1).

**Table 4:** Stroke SMART simulation results of the true and estimated probability of success of each dynamic treatment regimen Caption: Performance of dynamic treatment regime when n=1500. The treatment effects for the 2.5 and 97.5^th^ percentiles of the arms of all simulated trials are represented by the 95% confidence interval. For example, in the scenario 1, for the first row representing receiving lytic A first, receiving usual care if an initial response occurs and if no response, receiving the new medication in stage 2 (instead of cath lab) only 2.5% of simulated trials returned overall good outcome probability of 0.58 or greater for that arm (with the given truth for that scenario equal to 0.53). A=Lytic A; B=Lytic B; r=responder; nr=non-responder; UC=usual care; NM=New Medicine; CL=Cath Lab

| Scenario/DTR | P(Success) |  |  |
| --- | --- | --- | --- |
|  | Truth | Estimated | 95% CI |
| Null |  |  |  |
| ArUCnrNM | 0.471 | 0.472 | 0.427, 0.517 |
| ArUCnrCL | 0.471 | 0.471 | 0.426, 0.517 |
| BrUCnrNM | 0.471 | 0.472 | 0.427, 0.517 |
| BrUCnrCL | 0.471 | 0.471 | 0.426, 0.516 |
| 1 |  |  |  |
| ArUCnrNM | 0.534 | 0.534 | 0.488, 0.580 |
| ArUCnrCL | 0.472 | 0.472 | 0.427, 0.518 |
| BrUCnrNM | 0.448 | 0.448 | 0.402, 0.494 |
| BrUCnrCL | 0.376 | 0.376 | 0.333, 0.422 |
| 2 |  |  |  |
| ArUCnrNM | 0.534 | 0.535 | 0.489, 0.580 |
| ArUCnrCL | 0.471 | 0.472 | 0.427, 0.517 |
| BrUCnrNM | 0.557 | 0.557 | 0.511, 0.602 |
| BrUCnrCL | 0.496 | 0.496 | 0.451, 0.541 |
| 3 |  |  |  |
| ArUCnrNM | 0.534 | 0.534 | 0.489, 0.580 |
| ArUCnrCL | 0.472 | 0.472 | 0.427, 0.517 |
| BrUCnrNM | 0.588 | 0.588 | 0.541, 0.632 |
| BrUCnrCL | 0.530 | 0.530 | 0.485, 0.575 |
| 4 |  |  |  |
| ArUCnrNM | 0.471 | 0.471 | 0.427, 0.517 |
| ArUCnrCL | 0.471 | 0.472 | 0.427, 0.518 |
| BrUCnrNM | 0.530 | 0.530 | 0.485, 0.575 |
| BrUCnrCL | 0.530 | 0.530 | 0.485, 0.575 |
| 5 |  |  |  |
| ArUCnrNM | 0.534 | 0.534 | 0.488, 0.580 |
| ArUCnrCL | 0.472 | 0.472 | 0.427, 0.517 |
| BrUCnrNM | 0.597 | 0.597 | 0.551, 0.641 |
| BrUCnrCL | 0.472 | 0.471 | 0.426, 0.517 |
| 6 |  |  |  |
| ArUCnrNM | 0.409 | 0.409 | 0.366, 0.453 |
| ArUCnrCL | 0.471 | 0.472 | 0.427, 0.517 |
| BrUCnrNM | 0.471 | 0.471 | 0.426, 0.516 |
| BrUCnrCL | 0.471 | 0.471 | 0.427, 0.517 |

In scenario 5, the comparison of DTRs that begin with A and B lytics followed by usual care shows the expected null result (both 0.472); however, for DTRs that give the new second stage treatment for non-responders, the outcome is numerically higher if the first treatment was lytic B – 0.597 (95% CI 0.551-0.641) versus if the first treatment was lytic A – 0.534 (95% CI 0.488–0.580).

In scenario 6, the comparison of DTRS that begin A and B followed by usual care is again null, however the use of the new second stage treatment following A demonstrates worse outcomes – 0.409 (95% CI 0.366-0.453).

As expected, the overall type I error is well controlled by the SMART (Table 5, null scenario), with the proportion of false positive trials in the null case approximately at 0.05 from sample sizes of 700 to 2000. In scenario 1, the power for the SMART (that one of the four DTRs is statistically different) is high across the sample sizes from 700-2000. For the 700 patient version of scenario 1, power is 0.80 to find a significant difference in the DTR versus 0.95 in the traditional trial to find only a significant different in lytics A and B. Considering the 1500 patient sized trials, the SMART has higher power for scenario 2 (0.766) when compared to the traditional trial (0.27). SMARTs are effective tools for constructing high quality DTRs and to find differences between DTRs. SMARTs can detect differences with greater power than a standard trial when, for example, the effect of the first-stage treatments was small, but there is a synergistic effect when a first-stage treatment is followed by a particular second stage treatment. This hypothesis (does a new medication work for the second stage versus usual care) would not have been tested in the traditional A versus B trial, but would have been evaluated separately in the futility trial.

**Table 5:** Number of individuals consistent with each dynamic treatment regimen (DTR) and the power Caption: Power indicates the ability to detect any difference in treatment effects across the four DTRs for each given scenario and sample size. A=1ytic A; B=1ytic B; r=responder; nr=non-responder; UC=usual care; NM=New Medicine; CL=Cath Lab

| Scenario | N |  |  |  |  | Power |
| --- | --- | --- | --- | --- | --- | --- |
|  | Total | ArUCnrNM | ArUCnrCL | BrUCnrNM | BrBUCnrCL |  |
| Null | 2000 | 686 | 686 | 686 | 686 | 0.049 |
|  | 1500 | 515 | 515 | 515 | 515 | 0.045 |
|  | 700 | 240 | 240 | 240 | 240 | 0.053 |
| 1 | 2000 | 686 | 686 | 641 | 641 | 1.0 |
|  | 1500 | 515 | 515 | 481 | 481 | 0.990 |
|  | 700 | 240 | 240 | 224 | 224 | 0.797 |
| 2 | 2000 | 687 | 687 | 696 | 695 | 0.876 |
|  | 1500 | 515 | 515 | 522 | 522 | 0.766 |
|  | 700 | 240 | 240 | 244 | 244 | 0.401 |
| 3 | 2000 | 686 | 686 | 710 | 709 | 0.961 |
|  | 1500 | 515 | 515 | 532 | 532 | 0.885 |
|  | 700 | 240 | 240 | 249 | 249 | 0.527 |
| 4 | 2000 | 686 | 686 | 709 | 708 | 0.590 |
|  | 1500 | 516 | 516 | 531 | 531 | 0.449 |
|  | 700 | 241 | 241 | 248 | 248 | 0.215 |
| 5 | 2000 | 687 | 686 | 686 | 686 | 1.0 |
|  | 1500 | 515 | 515 | 515 | 515 | 0.995 |
|  | 700 | 240 | 240 | 240 | 240 | 0.836 |
| 6 | 2000 | 686 | 686 | 686 | 687 | 0.714 |
|  | 1500 | 515 | 515 | 515 | 515 | 0.588 |
|  | 700 | 240 | 240 | 240 | 240 | 0.291 |

## Discussion

Using numerical simulation, we have demonstrated the utility of an alternative clinical trial design to test sequential treatments for acute stroke. For modest effect sizes, the SMART design provides comparable statistical power to conventional, separate clinical trials (one large phase III and one small futility design) testing alternative thrombolytic regimens and new treatments for reperfusion in the setting of non-response to initial thrombolytic therapy. One of the most important benefits of using the SMART strategy is the ability to find differences in DTRs or tailored sequences of treatments or treatment interactions (synergies or antagonisms), as opposed to treatments at a single stage. There is great biological plausibility that initial thrombolytic regimens other than alteplase may be more effective, however the incremental benefit is likely to be small and challenging to detect. The ability for an alternative initial thrombolytic regimen to improve the performance of second stage treatments (mechanical thrombectomy or alternative pharmacotherapies to safely enable reperfusion) would simply not be detected using the current trial development paradigm. Also important, if an alternative thrombolytic regimen displaced alteplase as the drug of choice, it is entirely conceivable that the benefit / risk profile of adjunctive mechanical therapies would be substantially altered as well, possibly with more adverse events and it would be extremely challenging to conduct randomized trials in this area. In essence, when a new thrombolytic regimen supplants alteplase for stroke, we will be treating patients based on the intuition that the newer regimen will be just as safe with thombectomy as opposed to objectively obtained randomized clinical trial data. Some comparisons to historical data will be possible, but will have important limitations.

Such a consolidated design also offers practical benefits. Clinical trials are costly to start up, complete, and disseminate the results. Particularly in the academic stroke world, which in the United States is largely consolidated with the StrokeNET as the platform for multi-center trials both in the late exploratory and confirmatory phase, clinical trial volunteers are a limited and precious resource. The ability to concurrently answer two important questions, (i.e. is a new lytic regimen better than the old, and is there a faster, easier, safe, effective pharmacological alternative to thrombectomy) and understand the interaction between the two stages of treatment presents a useful alternative pathway for the development of acute stroke treatments. Acute stroke care is dynamic at the bedside, but clinical trials in this space have typically focused only on the first decision. The SMART design presented here allows learning to occur at two important time points. This could also be applied to combined interventions at other time points, such as an alternative treatment up front changing the rehabilitation trajectory and making different restorative or compensatory methods for physical and occupational therapy more effective.

This work has several important limitations. First, we developed treatment response scenarios and treatment effects based on the responses seen in the NINDS tissue plasminogen activator trial. As the goal of our project was to demonstrate the utility of competing designs, it would be straightforward for other researchers to simulate other scenarios using the design and code we have developed for this. Second, the final analysis of the SMART clinical trial as we have simulated uses an omnibus test that one of the four DTRs is different. As can be observed with the overlapping confidence intervals for some of the treatment arms that are clearly superior to one another (based on the truth of the scenario that was simulated), using this exact approach may lead to important scientific and regulatory difficulties with interpretation. Still, additional design and simulation work that balances the needs of the clinical investigators with what is inferentially possible within plausible sample sizes could further refine this to ensure that the design satisfies regulators and the stroke community. Finally, developing clinical scenarios and simulations is time consuming and requires substantial effort and iterative communication between statisticians and clinicians.(9) Current funding opportunities and promotion and tenure requirements in academia do not adequately reflect and compensate for this time. Despite this, we invest a large amount of resources both financially and in the limited pool of available acute stroke patients conducting clinical trials. Developing modern, innovative, and flexible clinical trial designs that can maximize learning in this space should be facilitated by funding models or design competitions that bring better designs to the table for consideration in peer review.

In summary, the use of SMART designs to investigate effective and realistic dynamic treatment regimens are promising ways to accelerate the discovery of new, effective treatments for stroke. The current treatment development pathway in academia has important shortcomings including lack of direct insights into how new combined treatments will work in the future. More attention to clinical trial design and simulation is warranted and will help achieve the goals of reducing stroke disability.

## Acknowledgments / Sources of Funding / Conflicts of Interest

The authors have no relevant conflicts of interest to report. The authors report relevant funding from the University of Michigan Cancer Center Biostatistics Training Grant 5T32CA083654 and the National Institutes of Neurological Disorders and Stroke Clinical Trials Methodology Course R25 NS088248.

## Supplementary Material

**Supplemental Table 1:**
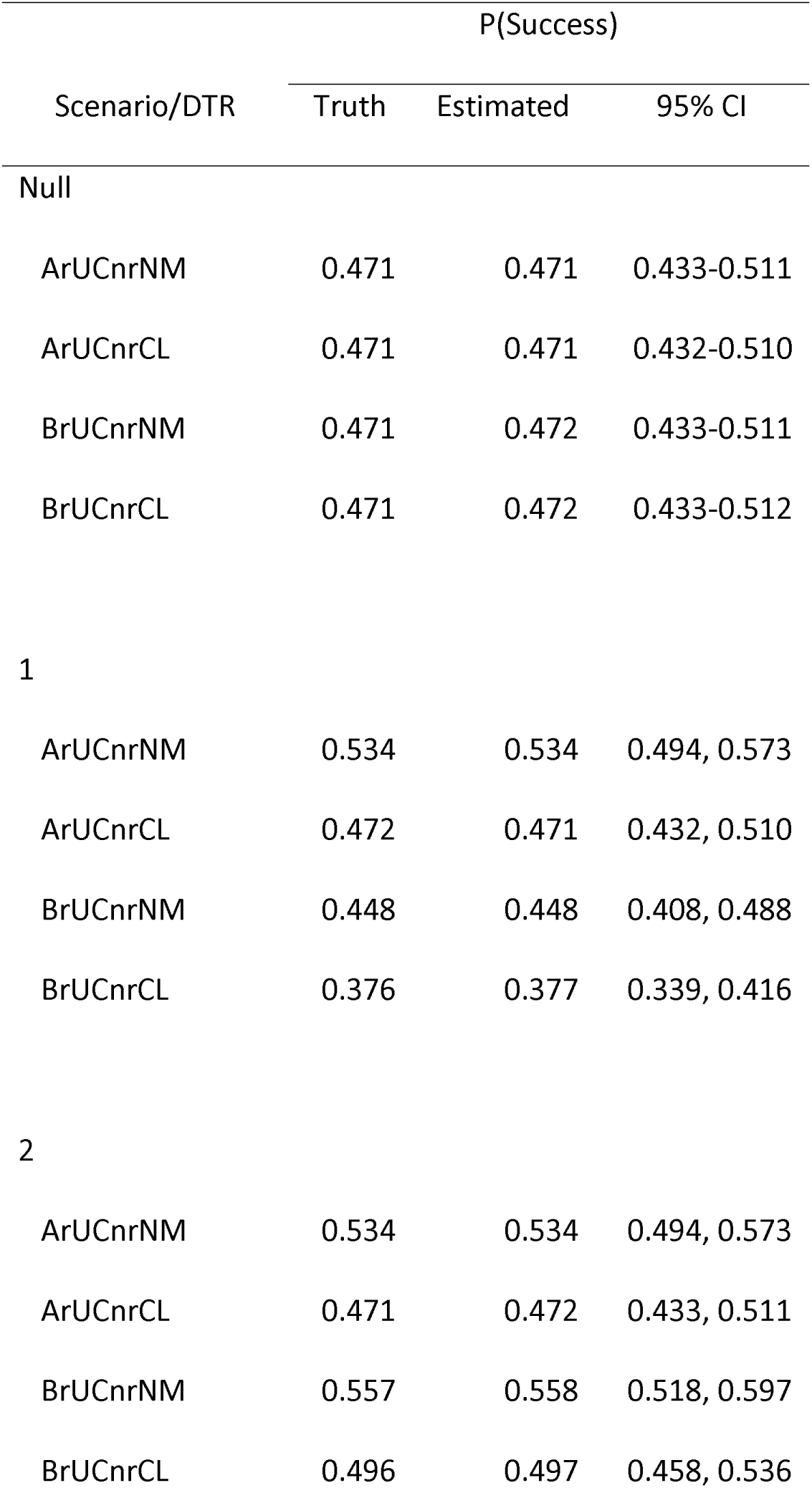
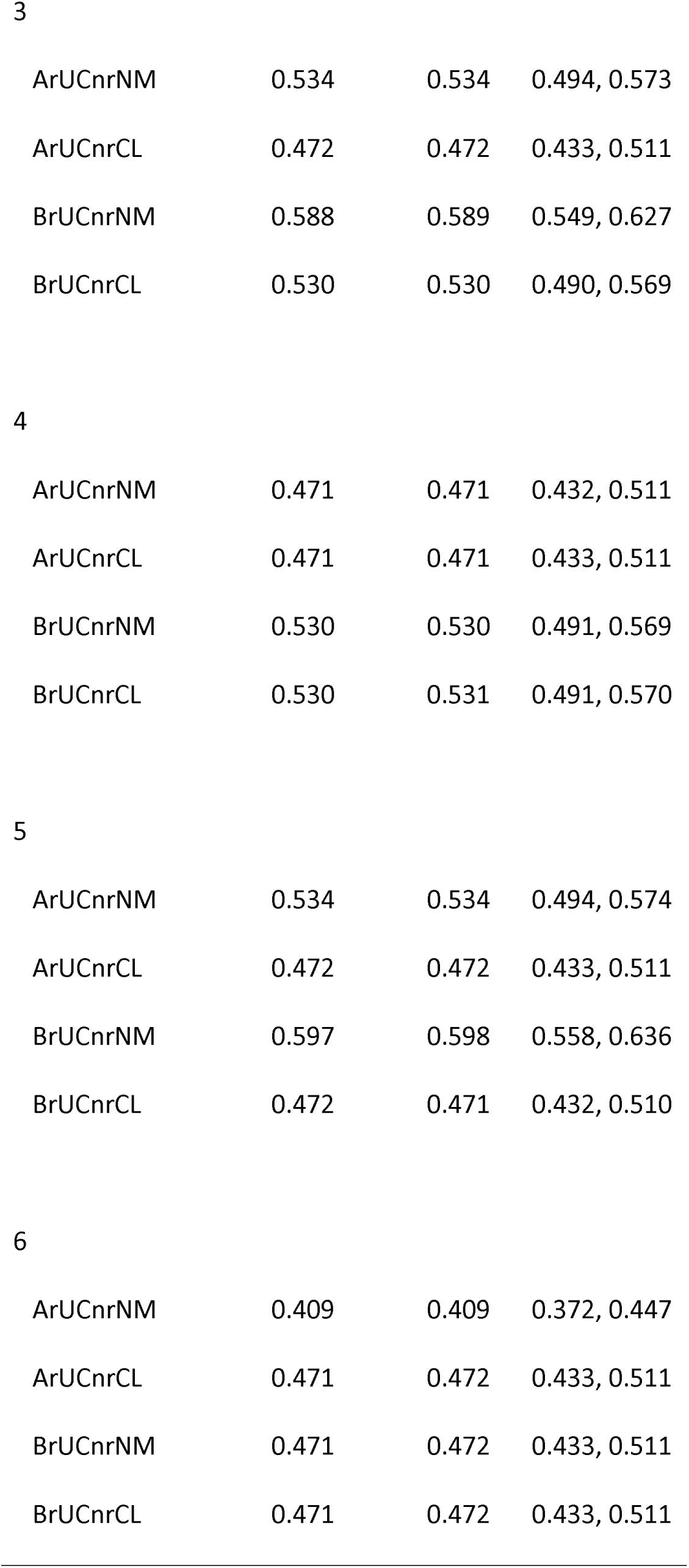
Stroke SMART simulation results of the true and estimate success of the dynamic treatment regimen Captions: Given sample size is 2000 for this table. A=1ytic A; B=1ytic B; r=responder; nr=non-responder; UC=usual care; NM=New Medicine; CL=Cath Lab

~~~
R code to implement SMART simulations
## DESCRIPTION OF VARIABLES:
## n: Sample Size
## qAr: Probability of Y=1 for those who are given first-stage treatment A and respond
## qBr: Probability of Y=1 for those who are given first-stage treatment B and respond
## qAnrWW: Probability of Y=1 for those who are given A, don't respond, and are assigned to WW
## qAnrCL: Probability of Y=1 for those who are given A, don't respond, and are assigned to CL
## qBnrWW: Probability of Y=1 for those who are given A, don't respond, and are assigned to WW
## qBnrCL: Probability of Y=1 for those who are given A, don't respond, and are assigned to CL
## prBI: Probability of response given B and low baseline
## prBh: Probability of response given B and high baseline
## alpha: Probability of Type-1 Error, defaults to 0.05
## niter: Number of iterations to carry out, defaults to 5000.
#####################################################################################
~~~

~~~
libra ry(aod)
libra ry(geepack)
libra ry(doBy)
#set seed
RNGkind("Mersenne-Twister")
set.seed(1111)
~~~

~~~
simulate.strokeSMART<-function(n,qAr,qBr,qAnrWW,qAnrCL,qBnrWW,qBnrCL,prBl, prBh, alpha=0.05,niter=5000){
~~~

~~~
N = n*niter
~~~

~~~
# model baseline NIHSS based on proportion data from NINDS
#create indicator for baseline: 1 if < 8, 0 otherwise
nihss. baseline.dichot<-rbinom(N, 1,115/509)
~~~

~~~
#create outcome probability vectors for each embedded DTR
pArAnrWW <-rep(NA,N)
pArAnrCL <-rep(NA,N)
pBrBnrWW <-rep(NA,N)
pBrBnrCL <-rep(NA,N)
pArAnrWW[nihss.baseline.dichot==1]<-(0.93)*qAr + (0.07)*qAnrWW
pArAnrCL[nihss.baseline.dichot==1]<-(0.93)*qAr + (0.07)*qAnrCL
pArAnrWW[nihss.baseline.dichot==0]<-(0.21)*qAr + (0.79)*qAnrWW
pArAnrCL[nihss. baseline. dichot==0]<-(0.21)*qAr + (0.79)*qAnrCL
pBrBnrWW[nihss.baseline.dichot==1]<-prBI*qBr + (1-prBI)*qBnrWW ##
pBrBnrCL[nihss.baseline.dichot==1]<-prBI*qBr + (1-prBI)*qBnrCL ##
pBrBnrWW[nihss.baseline.dichot==0]<-prBh*qBr + (1-prBh)*qBnrWW ##
pBrBnrCL[nihss.baseline.dichot==0]<-prBh*qBr + (1-prBh)*qBnrCL ##
~~~

~~~
#Miscellaneous initializations
omnibusCount=0
pairwiseCount=0
pairwiseCountWW=0
#percentDone = 0
~~~

~~~
#assign first-stage treatment
# 1 is A, -1 is B
stage1trt<-rbinom(N,1,.5)
stage1trtfstage1trt==0]<--1
~~~

~~~
#set response probability for A/B based on baseline NIHSS and simulate response
stage1resp<-rep(NA,N)
stage1respfnihss.baseline.dichot==1 & stage1trt==1]<-rbinom(sum(nihss.baseline.dichot==1 & stage1trt==1), 1,0.93)
stage1respfnihss.baseline.dichot==0 & stage1trt==1]<-rbinom(sum(nihss.baseline.dichot==0 & stage1trt==1),1,0.21)
stage1respfnihss.baseline.dichot==1 & stage1tr==1]<-rbinom(sum(nihss.baseline.dichot—1 & stage1trt==-1),1,prBl)
stage1respfnihss.baseline.dichot==0 & stage1trt==1]<-rbinom(sum(nihss.baseline.dichot==0 & stage1trt==-1),1,prBh)
~~~

~~~
#assign second-stage treatment to non-responders
# 1 is CL, -1 is WW
num.resp<-sum(stage1resp==1)
stage2t rt<- re p( N A, N )
stage2trt[stage1resp==0]<-rbinom(N-num.resp,1,.5)
stage2trt[stage2trt==0]<--1
~~~

~~~
#assign weights (2 per randomization)
weight<-rep(NA,N)
weight[stage1resp==1]<-2
weight[stage1resp==0]<-4
~~~

~~~
#simulate outcome Y
Y<-rep(NA,N)
Y[stage1trt==1 & stage1resp==1]<-rbinom(sum(stage1trt==1 & stage1resp==1),1,qAr)
Y[stage1trt==-1 & stage1resp==1]<-rbinom(sum(stage1trt==-1 & stage1resp==1),1,qBr)
Y[stage1trt==1 & stage1resp==0 & stage2trt==1]<-rbinom(sum(stage1trt==1 & stage1resp==0 & stage2trt==1),1,qAnrCL)
Y[stage1trt==1 & stage1resp==0 & stage2trt==1]<-rbinom(sum(stage1trt==1 & stage1resp==0 & stage2trt==-1),1,qAnrWW)
Y[stage1trt==-1 & stage1resp==0 & stage2trt==1]<-rbinom(sum(stage1trt==-1 & stage1resp==0 & stage2trt==1),1,qBnrCL)
Y[stage1trt==-1 & stage1resp==0 & stage2trt==1]<-rbinom(sum(stage1trt==-1 & stage1resp==0 & stage2trt==-1),1,qBnrWW)
#identify DTR (path-specific, e.g., Ar, AnrWW, and AnrCL are considered separately)
# 11 is Ar, 100 is AnrWW, 101 is AnrCL, -11 is Br, -100 is BnrWW, -101 is BnrCL
DTR<-rep(NA,N)
DTR[stage1trt==1 & stage1resp==1]<-11 #Ar
DTR[stage1trt==1 & stage1resp==1]<--11 #Br
DTR[stage1trt==1 & stage1resp==0 & stage2trt==1]<-101 #AnrCL
DTR[stage1trt==1 & stage1resp==0 & stage2trt==-1]<-100 #AnrWW
DTR[stage1trt==1 & stage1resp==0 & stage2trt==1]<--101 #BnrCL
DTR[stage1trt==1 & stage1resp==0 & stage2trt==1]<--100 #BnrWW
~~~

~~~
#compute mean success probabilities
#Note: 0.309 is overall response probability
mspArAnrCL = (sum(Y[stage1trt==1 & stage1resp==1])/sum(DTR==11))*0.309 + (sum(Y[stage1trt==1 & stage1resp==0 & stage2trt==1])/sum(DTR==101))*(1-.309)
mspArAnrWW = (sum(Y[stage1trt==1 & stage1resp==1])/sum(DTR==11))*0.309 + (sum(Y[stage1trt==1 & stage1resp==0 & stage2trt==-1])/sum(DTR==100))*(1-.309)
mspBrBnrCL = (sum(Y[stage1trt==-1 & stage1resp==1])/sum(DTR==11))*0.309 + (sum(Y[stage1trt==1 & stage1resp==0 & stage2trt==1])/sum(DTR==-101))*(1-.309)
mspBrBnrWW = (sum(Y[stage1trt==-1 & stage1resp==1])/sum(DTR==11))*0.309 + (sum(Y[stage1trt==-1 & stage1resp==0 & stage2trt==-1])/sum(DTR==-100))*(1-.309)
~~~

~~~
#create bigdata matrix from which to sample.
bigdata<-matrix(c(stage1trt,stage1resp,stage2trt,Y, weight),ncol=5)
colnames(bigdata)<-c("stage1trt","stage1resp","stage2trt","Y", "weight")
chooserows<-matrix(c(sample(1:N,N,replace=T)),nrow=niter,ncobn) #get random sample of niter rows for each of the n subjects
data<-matrix(nrow=n,ncoN5)
~~~

~~~
#initialize probability and count matrices
DTRcount<-matrix(nrow=niter,ncoN4)
DTRIogitProb<-matrix(nrow=niter,ncoN4)
DTRIogitVar<-matrix(nrow=niter,ncoN4)
DTRIogitConf<-matrix(nrow=niter,ncoN8)
colnames(DTRcount)<-c("ACL","AWW","BCL","BWW")
colnames(DTRlogitProb)<-c("ACL","AWW","BCL","BWW")
colnames(DTRlogitVar)<-c("ACL","AWW","BCL","BWW")
colnames(DTRlogitConf)<-c("ACL Lower", "AWW Lower", "BCL Lower", "BWW Lower", "ACL Upper", "AWW Upper", "BCL Upper", "BWW Upper")
looptimeSTART<-proc.time()[3]
for (i in seq(from=1,to=niter,by=1)){
#create data matrix
data<-bigdata[chooserows[1,],] #consider only n observations
lengthNR<-dim(data[data[,2]==0,])[1] #number of non-responders to first-stage treatment
dataNR<-cbind(data[data[,2]==0,], c(seq(1,lengthNR,by=1))) #create dataset of just nonresponders to stage1trt and assign an ID
dataresp1<-data[data[,2]==1,] #subsetset of patients who responded to first-stage treatment
dataresp1[,3]<-1 #assign responders to stage2trt=1
lengthR<-dim(dataresp1)[1] #number of responders
dataresp1<-cbind(dataresp1, c(seq(lengthNR+1,lengthNR+lengthR))) #assign ID to responders
dataresp2<-data[data[,2]==1,] #subset of patients who responded to first-stage treatment
dataresp2[,3]<--1 #assign responders to stage2trt=-1
dataresp2<-cbind(dataresp2, c(seq(lengthNR+1,lengthNR+lengthR))) #assign ID
~~~

~~~
#Count individuals consistent with each DTR
DTRcount[i,1]<- sum(data[,1]==1 & data[,2]==1) + sum(data[,1]==1 & data[,2]==0 & data[,3]==1) #ArAnrCL
DTRcount[i,2]<- sum(data[,1]==1 & data[,2]==1) + sum(data[,1]==1 & data[,2]==0 & data[,3]==-1) #ArAnrWW
DTRcount[i,3]<- sum(data[,1]==-1 & data[,2]==1) + sum(data[,1]==-1 & data[,2]==0 & data[,3]==1) #BrBnrCL
DTRcount[i,4]<- sum(data[,1]==-1 & data[,2]==1) + sum(data[,1]==-1 & data[,2]==0 & data[,3]==-1) #BrBnrWW
~~~

~~~
#Replicate data
repdata<-rbind(dataNR, dataresp1, dataresp2)
colnames(repdata)<-c("stage1trt","response","stage2trt","Y","weight", "id")
inter<-c(repdata[,1]*repdata[,3]) #"interaction" variable
repdata<-cbind( re pdata, inter)
repdata<-as.data.frame(repdata)
r<-sum(repdata$response==1)/2/n
sortrepdata<-repdata[order(repdata$id), ]
~~~

~~~
#Estmate full and null models
glmrepdata3<-geeglm(Y ~ stage1trt+stage2trt+inter, id=id, weights=weight, family = binomial("logit"),corstr-'independence",data=sortrepdata)
glmrepdata3RED<-geeglm(Y ~1, id=id, weights=weight, family = binomial("logit"),corstr-'independence",data=sortrepdata)
#summary(glmrepdata3)
~~~

~~~
#Omnibus test
omnibustest<-anova(glmrepdata3,glmrepdata3RED, test-'Chisq")
if (omnibustest$'P(> | Chi | )'<alpha)
omnibusCount = omnibusCount + 1
~~~

~~~
#Pairwise test: ArAnrCL vs. BrBnrCL
lpairACLBCL<-cbind(0,2,0,2)
pairACLBCL<-wald.test(b = coef(glmrepdata3), Sigma = vcov(glmrepdata3), L=1pairACLBCL)$result$chi2[3]
if (pairACLBCL<alpha)
#Pairwise test: ArAnrWW vs. BrBnrWW
lpairAWWBWW<-cbind(0,2,0,-2)
pairAWWBWW<-wald.test(b = coef(glmrepdata3), Sigma = vcov(glmrepdata3), L=1pairAWWBWW)$result$chi2[3]
if (pairAWWBWW<alpha)
pairwiseCountWW = pairwiseCountWW + 1
~~~

~~~
#Estimate and store logit(DTRprob) and confidence interval for each DTR
#1,1,1,1<=>ArAnrCL
# 1,1,-1,-1 <=> ArAnrWW
# 1,-1,1,-1 <=> BrBnrCL
# 1,-1,-1,1 <=> BrBnrWW
est<-esticon(glmrepdata3,matrix(c(1,1,1,1,1,1,-1,-1,1,-1,1,-1,1,-1,-1,1),ncol=4))
DTRlogitProb[i,]<-est$Estimate
DTRIogitVar[i,]<-(est$Std.Error)^^^2
DTRIogitConf[i,]<-c(est$Lower,est$Upper)
~~~

~~~
#crude iteration indicator (for progress purposes)
#if (i%%50 == 0){
# percentDone = percentDone + 1
# cat(percentDone,'% completed.','\n')
}
looptimeEND<-proc.time()[3]
~~~

~~~
powerOmnibus<-omnibusCount/niter
powerPairwiseACLBCL<-pairwiseCount/niter
powerPairwiseAWWBWWc-pairwiseCountWW/niter
~~~

~~~
#Compute expit probability and Cl's
DTRprob<-exp(DTRlogitProb)/(1+exp(DTRlogitProb))
DTRconf<-exp(DTRlogitConf)/(1+exp(DTRIogitConf))
~~~

~~~
#Compute variances for each estimated DTR probability (columns of DTRprob)
varArAnrCL<-var(DTRprob[,1])
varArAnrWW<-var(DTRprob[,2])
varBrBnrCL<-var(DTRprob[,3])
varBrBnrWW<-var(DTRprob[,4])
~~~

~~~
#compute empirical confidence interval – method 1, using CLT
empDTRconf1<-matrix(NA,nrow=4,ncol=2)
empDTRconf1[1,]<-c(mean(DTRprob[,1])-
1.96*sqrt(varArAnrCL),mean(DTRprob[,1])+1.96*sqrt(varArAnrCL))
    empDTRconf1[2,]<-c(mean(DTRprob[,2])-
1.96*sqrt(varArAnrWW),mean(DTRprob[,2])+1.96*sqrt(varArAnrWW))
    empDTRconf1[3,]<-c(mean(DTRprob[,3])-
1.96*sqrt(varBrBnrCL),mean(DTRprob[,3])+1.96*sqrt(varBrBnrCL))
    empDTRconf1[4,]<-c(mean(DTRprob[,4])-
1.96*sqrt(varBrBnrWW),mean(DTRprob[,4])+1.96*sqrt(varBrBnrWW))
~~~

~~~
#compute empirical confidence interval - method 2, using percentiles
empDTRconf2<-matrix(NA,nrow=4,ncol=2)
empDTRconf2[1,]<-quantile(DTRprob[,1],c(.025,.975),na.rm=TRUE) #ArAnrCL
empDTRconf2[2,]<-quantile(DTRprob[,2],c(.025,.975),na.rm=TRUE) #ArAnrWW
empDTRconf2[3,]<-quantile(DTRprob[,3],c(.025,.975),na.rm=TRUE) #BrBnrCL
empDTRconf2[4,]<-quantile(DTRprob[,4],c(.025,.975),na.rm=TRUE) #BrBnrWW
~~~

~~~
#t = time()-t0
#print(t)
~~~

~~~
cat(’ ~~~~~~~~~~~~~~~~~~~~~~~~','\n',
‘POWER with’ n, ‘subjects:’, '\n',\
‘of the Omnibus Test:’,round(powerOmnibus,4), '\n',
‘of the ACLBCL Pairwise Test:’, round(powerPairwiseACLBCL,4),'\n',
‘of the AWWBWW Pairwise Test:,round(powerPairwiseAWWBWW,4),'\n',
‘~~~~~~~~~~~~~~~~~~~~~~~~~~~’, '\n',\
cat (‘~~~~~~~~~~~~~~~~~~~~~~~; '\n',\
‘ESTIMATED success probability per DTR:', '\n',
' ArAnrCL: ',mean(DTRprob[,1]),'\n',
' ArAnrWW: ',mean(DTRprob[,2]),'\n',
' BrBnrCL: ',mean(DTRprob[,3]),'\n',
' BrBnrWW: ',mean(DTRprob[,4]),'\n', _An_,_}_
‘~~~~~~~~~~~~~~~~~~~~~~~~~~~’, '\n',\
cat (‘~~~~~~~~~~~~~~~~~~~~~~~; '\n',\
' ESTIMATED 95% confidence interval for P(Success)\n',
' ArAnrCL: (',mean(DTRconf[,1]),’,’,mean(DTRconf[,5]),') \n',
' ArAnrWW: (',mean(DTRconf[,2]),’,’,mean(DTRconf[,6]),') \n',
' BrBnrCL: (',mean(DTRconf[,3]),’,’,mean(DTRconf[,7]),') \n',
' BrBnrWW: (',mean(DTRconf[,4]),’,’,mean(DTRconf[,8]),') ‘\n',
‘~~~~~~~~~~~~~~~~~~~~~~~~~~~’, '\n',\
cat (‘~~~~~~~~~~~~~~~~~~~~~~~; '\n',\
' EMPIRICAL 95% confidence interval for P(Success) (via variance)\n',
' ArAnrCL: C,empDTRconfl[11],',',empDTRconf1[1,2],') \n',
' ArAnrWW: C,empDTRconfl[2,1]',',,empDTRconfl[2,2],') \n',
' BrBnrCL: C,empDTRconfl[3,1],',',empDTRconf1[3,2],') \n',
' BrBnrWW: (\empDTRconfl[4,1],',',empDTRconf1[4,2],') \n\
‘~~~~~~~~~~~~~~~~~~~~~~~~~~~’, '\n',\
cat (‘~~~~~~~~~~~~~~~~~~~~~~~; '\n',\
'EMPIRICAL 95% confidence interval for P(Success) (via percentile)\n',
'ArAnrCL: C,empDTRconf2[1,1],7,empDTRconf2[1,2],') \n',
' ArAnrWW: C,empDTRconf2[2,1],',',empDTRconf2[2,2],') \n',
' BrBnrCL: C,empDTRconf2[3,1],',',empDTRconf2[3,2],') \n',
' BrBnrWW: (',empDTRconf2[4,1],',',empDTRconf2[4,2],') \n',
‘~~~~~~~~~~~~~~~~~~~~~~~~~~~’, '\n',\
cat (‘~~~~~~~~~~~~~~~~~~~~~~~; '\n',\
' Mean number of individuals per DTR: \n',
' ArAnrCL: ',mean(DTRcount[,1]),'\n',
' ArAnrWW: ',mean(DTRcount[,2]),'\n',
' BrBnrCL: ',mean(DTRcount[,3]),'\n',
' BrBnrWW: ',mean(DTRcount[,4]),'\n',
‘~~~~~~~~~~~~~~~~~~~~~~~~~~~’, '\n',\
cat (‘~~~~~~~~~~~~~~~~~~~~~~~; '\n',\
' TRUE success probability per DTR:', '\n',
' ArAnrCL: ',mean(pArAnrCL),'\n',
' ArAnrWW: ',mean(pArAnrWW),'\n',
' BrBnrCL: ',mean(pBrBnrCL),'\n',
' BrBnrWW: ',mean(pBrBnrWW),'\n',
‘~~~~~~~~~~~~~~~~~~~~~~~~~~~’, '\n',\
cat (‘~~~~~~~~~~~~~~~~~~~~~~~; '\n',\
'Time Spent on Loop:', looptimeEND-looptimeSTART, '\n',
}
cat('NULL SCENARIO: \n')
simulate.strokeSMART(2000,0.76,0.76,0.3,0.3,0.3,0.3,0.93,0.21, alpha=0.05,niter=5000)
~~~

